# Fleeting Effects of Incentives: Adult Age Differences in ERP Measures of Motivated Attention

**DOI:** 10.1101/2022.04.18.488665

**Authors:** Margot D. Sullivan, Farrah Kudus, Benjamin J. Dyson, Julia Spaniol

## Abstract

Reward-based motivation is associated with transient and sustained dopaminergic activity and with modulatory effects on attention and cognitive control. Age-related changes in the dopamine system are well documented, but little is known about age differences in the temporal dynamics of motivational influences on cognitive functions. The current study examined the effects of financial incentives on visual attention using event-related potentials (ERPs). Participants (26 younger, aged 18-33; 24 older, aged 65-95) completed an incentivized flanker task in which trial-level incentive cues signaled the availability of performance-contingent reward, and subsequent alerting cues signaled the onset of the flanker target. ERP components of interest included incentive-cue P2 and CNV, alerting-cue N1, target N1, and target P3. Transient effects of incentives were assessed by comparing ERP amplitudes across incentive and non-incentive trials from mixed-incentive blocks. Younger adults showed transient effects of incentives on all components, whereas older adults showed effects for incentive-cue P2 and alerting-cue N1 only. Sustained effects of incentives were assessed by comparing ERP amplitudes across non-incentive trials from mixed-incentive blocks and non-incentive trials from pure non-incentive blocks. Both age groups showed sustained effects of incentives on cue-locked ERPs, but only younger adults showed sustained effects on target-locked ERPs. Reaction-time patterns mirrored the ERP findings, in that younger adults showed greater incentive-based modulation than older adults. Overall, these findings suggest that both transient and sustained effects of incentives on visual attention are more fleeting for older than younger adults, consistent with widespread alterations in dopaminergic neuromodulation in aging.

## Introduction

Age-related declines in attention and cognitive control are consequential for day-to-day complex tasks, such as maintaining attention and avoiding interference while driving (Adrian, Moessinger, Charles, & Postal, 2019). If incentives can effectively modulate attention and cognitive control in older adults, this would have practical implications for improving these abilities in later life. However, the underlying neural mechanisms and time course of how incentives influence attention and cognitive control in younger and older adults remains poorly understood (Ferdinand & Czernochowski, 2018). The current study aimed to identify age differences in the temporal dynamics of incentive-based modulation of visual attention in the context of an incentivized flanker task.

### Incentive effects on early vs. late components of attention and cognitive control

Behavioral and neural evidence suggests that incentives primarily influence proactive control processes (Botvinick & Braver, 2015; Hefer & Dreisbach, 2016; Notebaert & Braem, 2016). Proactive control emerges as early, anticipatory avoidance of upcoming interference, whereas reactive control occurs later in time and is involved in the detection and resolution of interference. Both processes are inherent to the dual-mechanisms-of-control theory (DMC; Braver & Barch, 2002; Braver, Paxton, Locke, & Barch, 2009; Braver, 2012). DMC theory suggests that age-related deficits in cognitive control are due to declines in dopamine and lateral prefrontal cortex system interactions. While certain studies indicate that older adults have preserved reactive control in the face of worse proactive control in relation to younger adults (Braver, 2012; Kopp, Lange, Howe, & Wessel, 2014; Paxton, Barch, Racine, & Braver, 2008), others have shown proactive control to be intact in older adults (Berger, Richards, & Davelaar, 2019; Staub, Doignon-Camus, Bacon, & Bonnefond, 2014), and thus, results may depend on task demands.

Chiew and Braver (2016) demonstrated that the availability of incentives improved proactive attentional control in younger adults performing the Eriksen flanker task (Eriksen & Eriksen, 1974), as participants were better able to use predictive task cues and showed a reduced interference cost on response speed. The timing of the incentive cue was crucial for performance, as when the incentive cue was presented in concert or in advance of tasks cues, incentives improved preparatory/proactive control processes, however when the incentive cue was presented after the predictive cue and closer in time to the task, incentives were associated with higher error rates. Similarly, in a task-switching paradigm, incentives were more beneficial to performance during a longer rather than shorter preparatory interval (Savine, Beck, Edwards, Chiew, & Braver, 2010). Nevertheless, reactive control processes may also benefit from incentives when reward signals align with the target without the presence of preparatory cues (see Krebs, Hopf, & Boehler, 2016, for review).

Examining event-related potentials (ERPs) in relation to proactive/preparatory vs. reactive processes allows for greater insight into the time course of incentive effects. The P2 is an early positive component peaking approximately 200ms post-stimulus, involved in automatic visual selective attention (Carretié, Hinojosa, Martín-Loeches, Mercado, & Tapia, 2004; Wastell & Kleinman, 1980), while the subsequent P3 component at 300ms is involved in a range of cognitive processes associated with target detection, including attention, memory, and context updating (Neuhaus et al., 2010; Polich, 2007). Schmitt, Ferdinand, and Kray (2015) examined the effects of financial incentives on control processes using ERPs and a modified AX continuous-performance task (AX-CPT). In this modified version an incentive cue preceded a context cue and both cues were presented in advance of the probe. In the context-dependent condition, the context cue influenced the response mapping to the subsequent probe, but not in the context-independent condition. Motivationally salient incentive cues (i.e., gains or losses) as opposed to neutral, non-incentive cues elicited larger P2 and P3 amplitudes in both groups at this initial stage of processing, although the P3 incentive effect was larger for younger than older adults, with other, more pronounced age differences emerging later in the trial. More specifically, at the time of the context cue, younger adults showed larger preparatory contingent negative variation (CNV -- a slow negative potential elicited at frontocentral sites that occurs between a warning cue and target onset; Galvao-Carmona et al., 2014) amplitude for losses, and also demonstrated greater reactive control later in the trial at the time of the probe in the form of larger N450 amplitude. In contrast, older adults showed larger context cue-locked P3 amplitude, but also an extended probe-locked P3, which was indicative of increased demands on working memory. These results support the idea that early rather than late attentional processes associated with incentives may be more stable with aging.

The Attention Network Test (ANT; Fan, McCandliss, Sommer, Raz, & Posner, 2002), which combines the flanker task (Eriksen & Eriksen, 1974) and the attentional cueing paradigm (Posner, 1980), separates attention into three distinct networks (i.e., alerting, orienting, and executive control) and has been used to explore age differences in incentive effects on attention. With aging, the orienting network is preserved, however, the effects of aging on the alerting and executive control networks are more mixed (Fernandez-Duque & Black, 2006; Gamboz, Zamarian, & Cavallero, 2010; Jennings, Dagenbach, Engle, & Funke, 2007). Behavioral work with younger and older adults has shown that the addition of an incentive condition to the ANT decreases overall response times (RTs), particularly for younger adults (Williams, Biel, Dyson, & Spaniol, 2017), with greater alerting effects in younger adults and greater flanker interference effects in older adults. However, faster responding by younger adults with incentives was associated with a cost to accuracy in the form of greater flanker interference. In this design, the incentive cue was present for the entire course of the trial and thus early vs. late incentive effects are confounded within the trial. However, these results clearly demonstrate that incentives enhance overall response times and alerting effects, mainly for younger adults, but also are associated with different strategy use by younger and older adults in terms of a speed-accuracy trade-off.

Williams et al. (2016) examined early and late ERP components in younger and older adults in relation to the ANT alerting cue and flanker target. Incentive information (i.e., gains vs. losses) was provided prior to each block, rather than at the trial-level. Posterior N1 amplitude, an early attentional component occurring between 150-250ms, was more negative for trials with a double cue vs. no cue (i.e., alerting effect) at both the cue and the target, with no differences between age groups. An alerting effect was also present for ANT cue-locked CNV amplitude, but age group differences did not emerge until the target. At the target, older adults, but not younger adults, demonstrated a typical flanker congruency effect, with reduced P3 amplitude for incongruent vs. congruent trials (Neuhaus et al., 2010; Galvao-Carmona et al., 2014), whereas only younger adults showed larger frontocentral N2 amplitude for incongruent vs. congruent targets. Thus, in this block-level incentive context, younger and older adults showed similar neural processing patterns for early, proactive/preparatory processes at the alerting cue, but not in terms of late, reactive processes at the target.

In summary, when incentive cues are presented ahead of upcoming cognitive tasks that require interference resolution, responses are faster, particularly in younger adults. Younger and older adults show similar patterns of neural activity at early stages of automatic attentional processes (P2 component) in relation to the incentive cue and at the alerting cue (posterior N1 component, CNV), but age differences emerge at later stages of processing at the incentive cue and the target (via P3 components). Despite similar alerting effects at the neural level between age groups, younger adults show greater alerting effects behaviorally. Younger adults show reduced flanker interference in their response times, but at a cost to accuracy, whereas older adults prioritize accuracy and also display greater congruency effects for P3 amplitude.

### Transient vs. sustained effects of incentives

On a broader timescale, incentives have been shown to elicit shifts in behavior (Marini, van den Berg, & Woldorff, 2015; Savine et al., 2010) and in the temporal dynamics of cognitive control via transient (i.e., trial-level) and sustained (i.e., block-level) effects (Jimura, Locke, & Braver, 2010; Chiew & Braver, 2013; Kostandyan et al., 2019). Transient effects of incentives are typically examined as changes in behavior or neural activity between incentive and non-incentive trials within reward blocks, whereas sustained effects of incentives represent differences between non-incentive trials embedded in reward blocks vs. neutral/baseline blocks (Chiew & Braver, 2013; Williams et al., 2018).

The distinction between transient and sustained effects of incentives mirrors two modes of dopamine transmission: rapid, transient firing vs. gradual, sustained activity (Chiew, Stanek, & Adcock, 2016; Niv, 2007; Yagishita, 2020). Transient dopamine release has been linked to the coding of predictable reward outcomes, whereas sustained dopamine release is associated more with reward uncertainty (Fiorillo, Tobler, & Schultz, 2003). Support for these distinct modes of release, and their relationship with reward anticipation, has been demonstrated in the memory domain with younger adults. Memory for images presented immediately after an incentive cue scaled with reward certainty, whereas images presented following a delay from the cue were best remembered when preceded by reward uncertainty (Stanek, Dickerson, Chiew, Clement, & Adcock, 2019).

In the domain of cognitive control, Chiew and Braver (2013) found that pupil diameter, an index of cognitive effort, increased in younger adults in both a transient and sustained manner during the AX-CPT, in which incentive cues were presented at the trial level prior to each contextual cue-probe pair. Transient effects occurred in advance of the probe, suggesting that incentives may enhance preparatory control. Similarly, Kostandyan et al. (2019) found that when incentive information was provided in a block-wise fashion vs. trial-wise during an incentivized flanker task, block-wise presentation resulted in sustained effects of incentives on pupil dilation, whereas trial-wise incentive cues elicited sustained effects as well as transient effects for pupil dilation along with a transient, speed-up effect for response times. Trial-wise presentation of incentive cues elicited larger P3 and CNV amplitude in relation to non-incentive cues, which again suggests that more immediate availability of incentive cues allows for greater proactive/preparatory control, with transient effects being more advantageous for performance.

Age-related alterations in the midbrain dopamine system are associated with changes in reward processing (Dreher, Meyer-Lindenberg, Kohn, & Berman, 2008) and cognition (Bäckman et al., 2000; Berry et al., 2016), but little is known about transient and sustained effects of incentives on cognitive control in older adults. Williams, Kudus, Dyson, and Spaniol (2018) examined ERP correlates of anticipatory and target processing in an incentivized flanker task that incorporated gains, losses, and neutral trials, in a mixed-block design with younger and older adults. Different patterns emerged for each age group during the earlier time window of the incentive cue-P3 component, with younger adults showing both transient and sustained effects of incentives, whereas incentive effects were limited to sustained effects of gains for older adults.

During late-stage anticipatory processing both age groups showed a similar pattern of transient incentive effects for CNV amplitude. At the level of the target, only older adults showed a congruency effect, with larger P3 amplitude for congruent vs. incongruent trials. Incentive effects were more prominent in younger adults, with both transient and sustained effects of incentives shown for target-P3 amplitude, whereas incentive effects were absent for older adults. Additionally, younger adults had larger cue- and target-locked P3 amplitude for incentive trials in relation to older adults. Incentives were associated with transient and sustained increases in response speed, but at a greater cost to accuracy for younger adults vs. older adults. Thus, while certain aspects of incentive effects were similar between age groups for anticipatory processing, at the stage of the target incentive effects have dissipated in older adults compared to younger adults.

In summary, trial-wise presentation of incentive cues within a mixed reward context elicits both transient and sustained effects on the temporal dynamics of cognitive control. Transient effects of incentives are conducive to preparatory processing and faster response speed, which is more detrimental to accuracy in younger adults. At the neural level, when older adult show incentive effects they tend to be present earlier within the preparatory phase of the trial, but are fleeting and disintegrate by the time of the target. In addition, for older adults, these effects are more sustained in nature such that they persist across trial types. In contrast, younger adults demonstrate more robust effects of incentives, showing transient and sustained effects during anticipatory processing and conflict resolution at the time of the target.

### The current study

Previous research has explored the relationship between incentives and alerting and between incentives and cognitive control. However, to our knowledge, no prior studies have investigated age differences in early and late stages of the neural time course of incentive processing for both alerting and cognitive control networks. The present study sought to extend previous behavioral (Williams et al., 2017) and neural findings (Williams et al., 2016; 2018) to characterize age differences in behavioral and neural responses at the level of the (1) incentive cue, (2) ANT alerting cue, and (3) target. Younger and older adult participants completed an incentivized flanker task, involving gain, loss, and neutral (i.e., non-incentivized) trials, with or without a preceding visual alerting cue, while ERPs were continuously recorded. Similar to prior studies investigating transient and sustained effects of incentives, motivational states were manipulated on a trial-by-trial basis through a mixed-block/event-related design (Chiew & Braver, 2013; Jimura et al., 2010; Williams et al., 2018).

Use of the temporal precision of ERPs enabled us to identify the stages at which incentive-based modulation of attention and cognitive control processes diverge between older and younger adults in terms of early vs. late occurring components, as well as the transient (i.e., trial-level) vs. sustained (block-level) nature of these effects. For these reasons, at the time of the incentive cue, we examined three components at frontocentral and centroparietal sites representing visual attention and preparatory processes: (1) P2 amplitude, (2) an early negative component amplitude^1^, and (3) later occurring CNV amplitude. At the time of the ANT cue, we examined posterior N1 amplitude, a negative early visual component associated with the alerting network. At the time of the target, we examined both posterior N1 amplitude, which has alsobeen shown to exhibit effects of alerting (Neuhaus et al., 2010; Galvao-Carmona et al., 2014), as well as frontocentral and centroparietal occurring target-P3 amplitude to investigate age-related processing differences in cognitive control.

Overall, we anticipated larger effects of incentives on neural processing and behavior in younger vs. older adults due to age-related changes in dopaminergic transmission. We specifically predicted that age differences in incentive-based modulation of attention and cognitive control should emerge at later stages of preparatory and target processing, rather than at earlier stages associated with more automatic attentional processes. We expected that incentive cue-locked P2 amplitude would increase with incentives and that posterior N1 amplitude time-locked to both the ANT cue and target would increase with alerting, and that both components should be largely age-invariant. CNV amplitude was expected to increase with incentives, however we had no specific predictions for age differences given mixed findings from prior studies (Schmitt, Ferdinand, & Kray, 2015; Williams et al., 2016, 2018). At the target, P3 amplitude was expected to be larger in younger adults vs. older adults. In line with Williams et al. (2016, 2018), older adults but not younger adults should show an effect of congruency on P3 amplitude. We predicted that younger adults would show transient and sustained effects of incentives on ERP components during anticipatory and target processing. Older adults were expected to show sustained effects of incentives during anticipatory processing that dissipate by the time of the target. We expected larger incentive-based modulation of behavior in younger vs. older adults, and greater prioritization of accurate performance by older adults. In addition, we explored whether incentives would affect the temporal dynamics of alerting.

## Method

### Participants

Twenty-six younger adults and 25 older adults participated in the current study. These group sample sizes were chosen to match those of prior studies in the literature (e.g., Williams et al., 2016; 2017; 2018). One older adult was excluded for having an accuracy level of < 60% in each experimental block. Included participants reported normal or corrected-to-normal hearing and vision, and were free of any major medical, neurological, or psychological problems.

Participants had to be native speakers of English or possess native-like English language proficiency, and have a minimum of 12 years of education. We aimed to recruit equal numbers of men and women in both age groups, and to match average years of education of both age groups. Group characteristics for this sample show typical age-related differences (see Table 1). All participants received $25 in cash in compensation for completing the study, which lasted 2-2.5 hours. Participants could win an additional bonus up to a maximum of $30 during the experimental task. We obtained approval for all study procedures from the Research Ethics Board of Ryerson University.

**Table 1.**
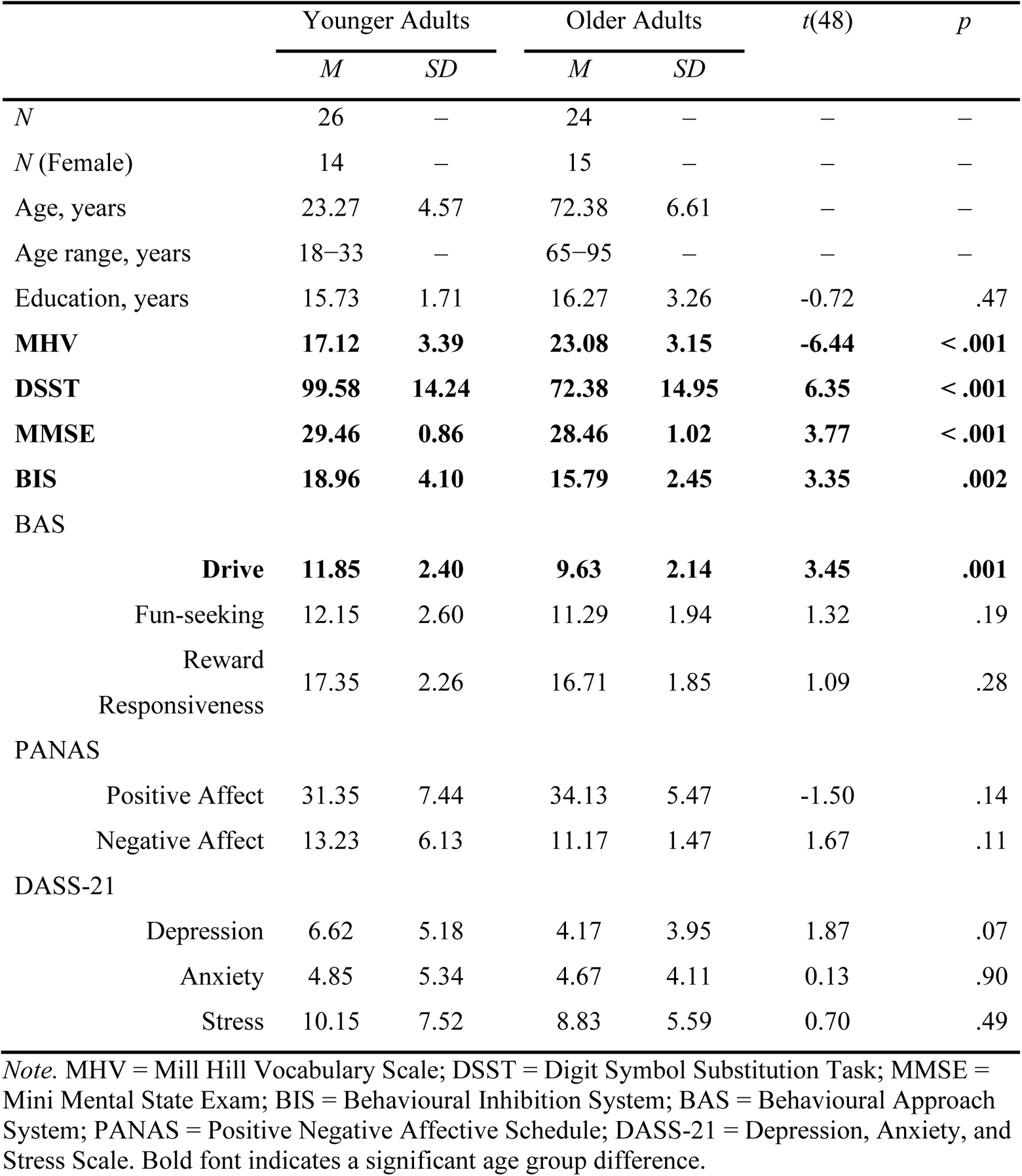
Group Characteristics

### Background Measures

Prior to the experiment, participants completed six background measures (see Table 1 for sample descriptives and inferential statistics). All participants completed a measure of cognitive functioning, the Mini Mental State Exam (MMSE; Folstein, Folstein, & McHugh, 1975), and scored at least 27 out of a possible 30. Participants also completed the Mill Hill Vocabulary Scale (MHV; Raven, 1982) to measure crystallized intelligence, the Digit Symbol Substitution Task (DSST; Wechsler, 1997) to measure perceptual-motor speed, and the Behavioural Inhibition/Behavioural Activation Scale (BIS/BAS; Carver & White, 1994) to measure dispositional sensitivity to reward and punishment. The BIS/BAS includes an inhibition scale and an activation scale, the latter of which comprises three subscales: drive, fun-seeking, and reward responsiveness. In addition, participants completed the Positive and Negative Affect Schedule (PANAS; Watson, Clark, & Tellegen, 1988) and the Depression Anxiety Stress Scales (DASS-21; Lovibond & Lovibond, 1995) to measure current affective state.

### Design and Apparatus

A modified version of the ANT was administered using Presentation software (Neurobehavioural Systems; Berkeley, CA), with participants seated approximately 50cm from the monitor. All stimuli were presented in white against a black background. Spatial and centre cues, which are typically used to estimate the efficiency of the orienting network in the ANT, were not included in this task. Furthermore, neutral flankers were not included based on prior evidence that ERPs elicited by congruent and neutral flankers are similar (Neuhaus et al., 2010). These modifications are analogous to those used in the ANT-G, a version which has previously been used with both healthy older adults and participants with mild cognitive impairment (Van Dam, 2013). The experimental design included the within-subjects factors of cue (no cue, double cue), congruency (congruent, incongruent), and block type: gain (G), loss (L), and neutral (N).

Within incentive blocks (i.e., gain and loss blocks), incentive availability varied trial-to-trial. On incentive trials (I), participants could win or lose 10 cents, whereas on neutral trials (N), no incentives were present. After taking into account the block-level and trial-level incentive manipulations, this design resulted in five unique trial types: gain-incentive (GI), gain-neutral (GN), loss-incentive (LI), loss-neutral (LN), and neutral-neutral (NN). Gain-neutral and loss-neutral trials were neutral trials presented in incentive blocks. In neutral blocks, only neutral trials were presented (NN). Lastly, the mixed-factorial design of this task also included age group (younger, older) as a between-subjects factor.

Within each of the five trial types, the four combinations of the two ANT cue conditions and two congruency conditions were presented with equal frequency. This resulted in 48 trials of each Trial Type x ANT Cue x Flanker combination. As a result of the neutral blocks having only a single trial type, neutral blocks included 48 trials each, whereas incentive blocks included 96 trials each. The total trial count over the course of the six experimental blocks was 480.

Two arrows flanked the target arrow on either side, pointing in either the same (congruent) or opposing (incongruent) direction as the central arrow. Within each condition, the central arrow pointed left or right, and the row of arrows appeared above or below the central fixation cross on 50% of the trials, respectively. On no-cue trials, no warning cue was presented, whereas on double-cue trials, two asterisks appeared on the screen above and below the central fixation point prior to the target onset. Lastly, feedback indicating whether the trial was successful (i.e., gain or non-loss) or unsuccessful (i.e., non-gain or loss) was presented after the participant’s response.

A schematic of the trial sequence is presented in Figure 1. Before each block, participants were informed whether the current block would be a neutral-only, gain, or loss block. At the start of each trial, an incentive cue appeared in the centre of the screen for 200ms (i.e., “&” on neutral trials, and “$” on incentive trials). Then, a fixation cross appeared for a variable duration that was randomly drawn from within the 400-1700ms interval. After this, the ANT cue (i.e., no cue or double cue) appeared for 100ms, followed by another fixation for 400ms. The target arrows then appeared above or below the fixation cross, and participants had up to 1,700ms to respond. After a response (or when the time limit had elapsed), the target disappeared and the fixation cross reappeared, followed by a feedback screen.

**Figure 1.**
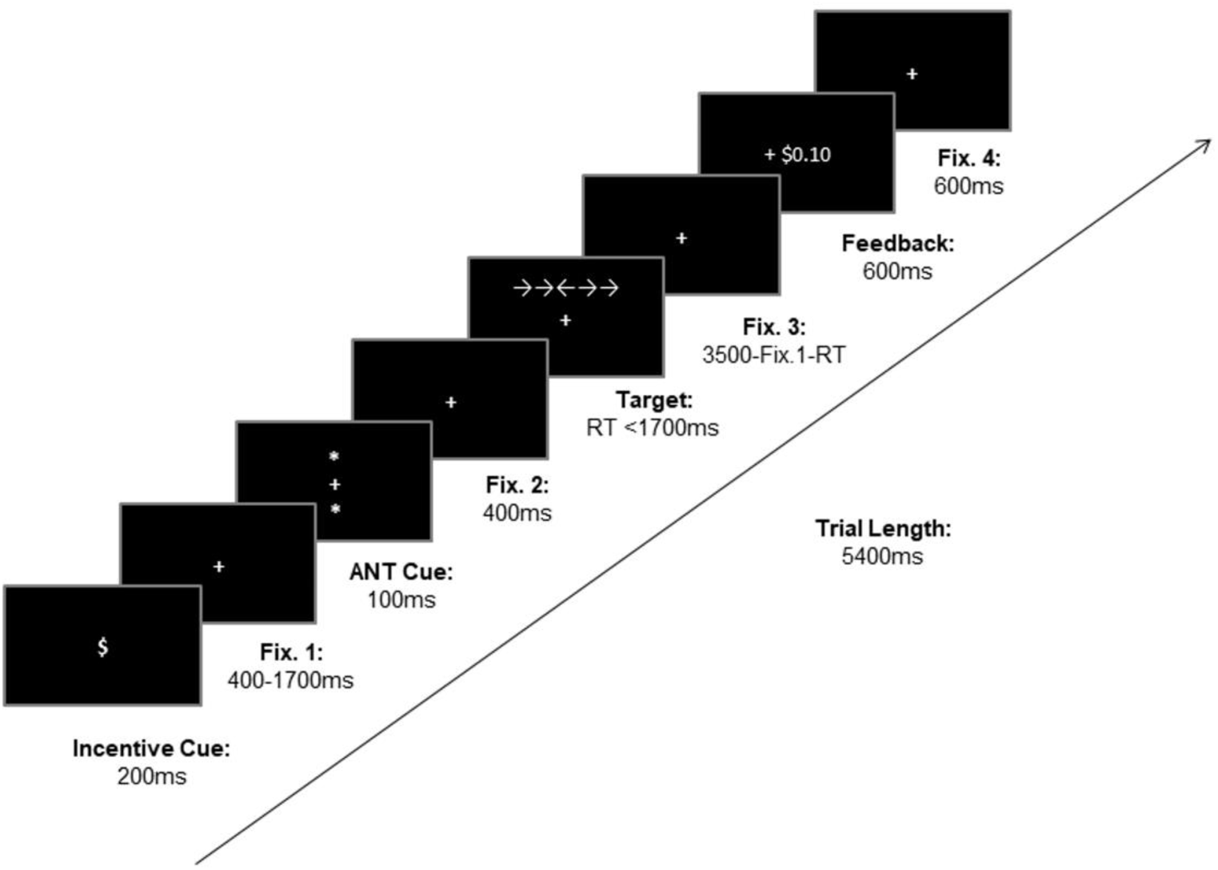
Trial Design for the Incentivized ANT, Showing a Single Gain-Incentive (GI) Trial with Positive Feedback

For GI trials, “+0.10” was presented for successful responses, whereas “+0.00” was presented for unsuccessful responses. For LI trials, “-0.00” and “-0.10” were presented for successful and unsuccessful responses, respectively. Feedback on neutral trials (GN, LN, and NN) was always the same (“#####”). The presentation of feedback was followed by a final fixation cross that stayed onscreen for 600ms, for a total trial length of 5,400ms. For a trial to be considered successful, participants were told that they had to be both accurate and faster than a time set by the computer. However, in reality, the task used a performance-adaptive response deadline (Williams et al., 2018), which ensured that each participant was successful on roughly 70% of incentivized trials. This was done by adjusting response time limits on a trial-by-trial basis. The response time limit was determined as the sum of cumulative average RT for each correct response (incentive trials only) plus an adjustable value. When the cumulative success rate was 70% or higher, the adjustable value was −10ms after correct responses and 0ms after incorrect or too-slow responses. When the cumulative success rate was below 70%, the adjustable value was +10ms after incorrect or too-slow responses and 0ms after correct responses. End-of-block feedback was also presented, which indicated the participant’s winnings (i.e., gains or avoided losses) during that block. No end-of-block feedback was given for neutral blocks.

### Procedure

After obtaining informed consent, participants completed the questionnaire measures and then moved on to the experimental task. Participants were instructed to respond to the direction (e.g., left or right) of the target arrow as quickly as possible, by pressing the “,” key with their right index finger if the central arrow pointed right, and the “X” key with their left index finger if the central arrow pointed left. Participants were told that they could earn a monetary bonus by giving fast and accurate responses. They completed three practice blocks of 24 trials each (1 neutral block, 1 gain block, 1 loss block), followed by the six experimental blocks. Between blocks, participants were required to take a break (at least 60s after the third block, at least 30s after all other blocks). The order of the experimental blocks was counterbalanced across participants within each age group. Following the task, participants were debriefed and received their compensation along with the bonus earned during the task. The bonus amount received was not significantly different for younger adults (*M* = 18.51, *SD* = 0.20) and older adults (*M =*18.49, *SD =* 0.20), *t*(48) *=* 0.35, *p* = .73.

### ERP Acquisition and Processing

Electrical brain activity was continuously collected for offline processing using an ActiveTwo system (BioSemi; Amsterdam, Netherlands) over an array of 64 electrodes, with a band-pass filter of 208 Hz and a 512 Hz sampling rate. Recordings were acquired from Ag/AgCl electrodes, which were connected to a cap (Cortech Solutions; Wilmington, NC) at 64 sites, according to the International 10-20 system. Six electrodes were attached externally, with two being placed on the right and left mastoids. Four electrodes were then used to record horizontal and vertical movements for both eyes by placing at the outer canthi and inferior orbits, respectively.

EEGlab (Delorme & Makeig, 2004) and ERPLab (Lopez-Calderon & Luck, 2014) were used to conduct off-line processing. Following the preprocessing procedure outlined by Williams et al. (2018), EEG data were referenced to the average of the right and left mastoids and were resampled as 256 Hz. High-pass (0.1 Hz, 12 dB/octave) and low-pass (30 Hz, 24 dB/octave) filters were applied to the continuous data. Then, cues and targets were epoched between 200ms pre-stimulus and 1000ms post-stimulus. Independent component analysis was used to correct artefacts (e.g., eye blinks, lateral eye movements). Finally, epochs containing values that exceeded a threshold of ±75 μV were automatically rejected. None of the participants had a total trial rejection rate greater than 30%.

### Behavioral Data Analysis

Blocks in which a participant failed to respond on more than 10% of trials, or in which accuracy was below 60% for any trial type, were excluded from further analyses. These criteria resulted in one older adult being excluded for having an accuracy rate below 60% for all blocks. To examine correct reaction time (RT) and accuracy effects across age groups and the different experimental conditions at the trial-level, we estimated multilevel models (MLMs) using the lme4 package in R (Bates et al., 2015). Main effects and interactions were tested using the Anova function from the car package in R (Fox & Weisberg, 2019) and are reported as Wald chi-square tests using type III sum of squares. Follow-up comparisons for significant effects and interactions (criterion, *p* < .05) were conducted using the R package emmeans (Lenth, 2020), using a false discovery rate *p*-value adjustment for multiple comparisons.

First, a multilevel linear regression (random intercept only) was estimated, with trials nested within subjects, regressing RT onto age group (younger, older), congruency (congruent, incongruent), ANT cue (no, double), and trial type collapsed across valence into three types (i.e., incentive trials [GI, LI], mixed-block neutral trials [GN, LN], and neutral trials [NN]). The ERP morphology was similar between GI and LI as well as between GN and LN trial types (see Figures 2–5) and preliminary behavioral analyses showed limited valence effects^2^. Therefore, we collapsed across valence to increase statistical power and to focus on our central aim of investigating the interaction between incentive and age group. In line with previous work (Fan et al., 2002; Mahoney et al., 2010), analyses of RT used correct responses only.

**Figure 2.**
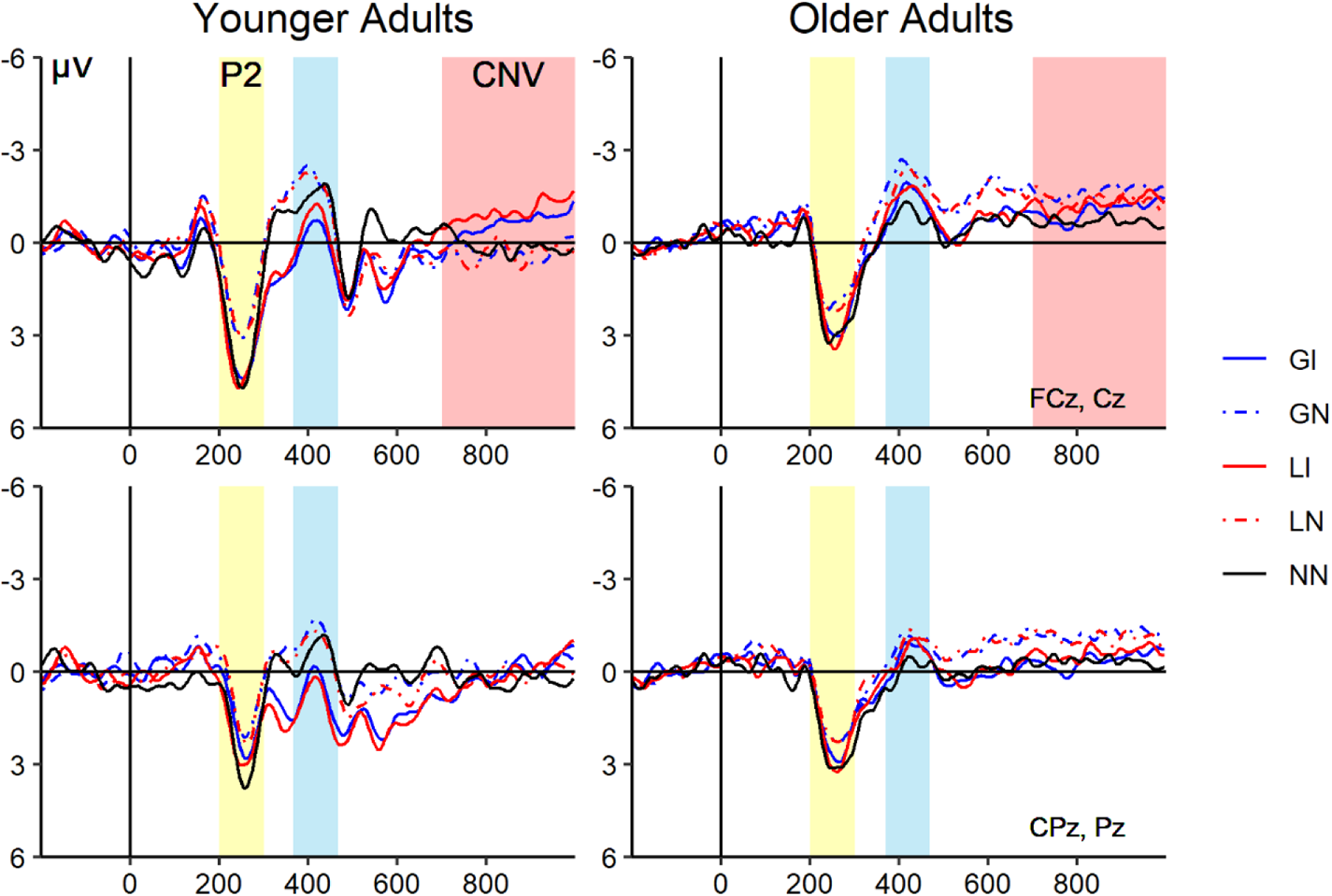
Incentive Cue-Locked ERPs by Age Group, Trial Type, and Region *Note.* ERP waveforms are averaged at frontocentral sites (top) and centroparietal sites (bottom). The yellow-shaded region corresponds to the time window for P2 amplitude, the blue-shaded region corresponds to the time window for the early negative component amplitude, and the pink-shaded region corresponds to the time window for CNV amplitude. GI = gain-incentive trials; GN = gain-neutral trials; LI = loss-incentive trials; LN = loss-neutral trials; NN = neutral-neutral trials.

**Figure 3.**
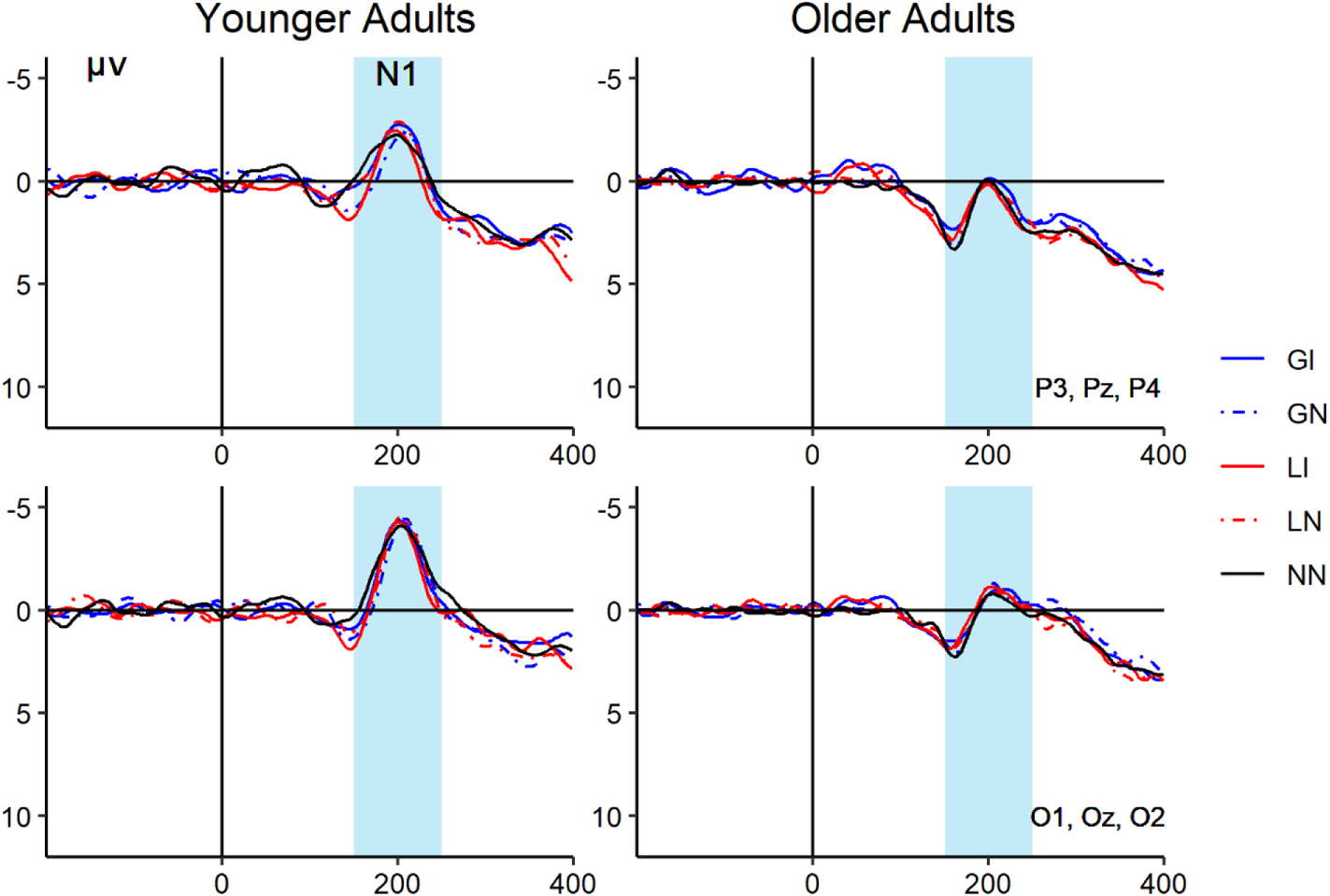
ANT Cue-Locked Difference Wave ERPs by Age Group, Trial Type, and Region *Note.* ERP difference waveforms are averaged at parietal sites (top) and occipital sites (bottom). Difference waveforms represent the double cue minus the no cue condition. The blue shaded region corresponds to the time window for posterior N1 amplitude.

**Figure 4.**
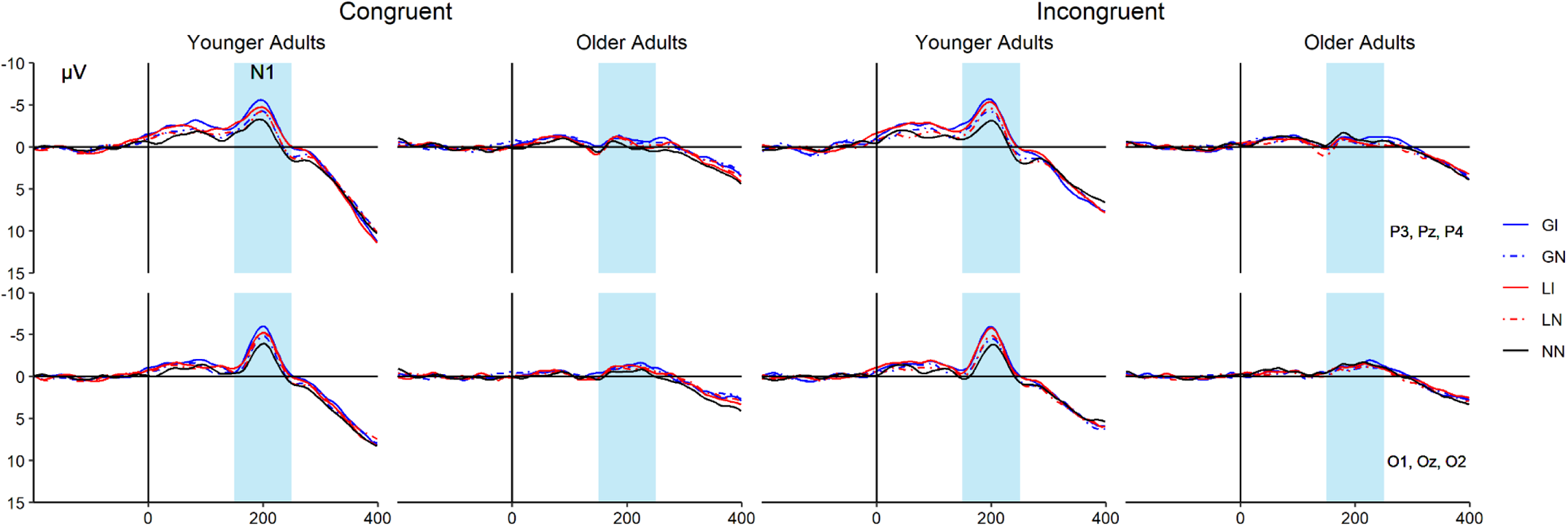
Target-Locked ERPs by Congruency, Age Group, Trial Type, and Region for Posterior N1 Amplitude *Note.* ERP waveforms are averaged at parietal sites (top) and occipital sites (bottom). The blue shaded region corresponds to the time window for posterior N1 amplitude. GI = gain-incentive trials; GN = gain-neutral trials; LI = loss-incentive trials; LN = loss-neutral trials; NN = neutral-neutral trials.

**Figure 5.**
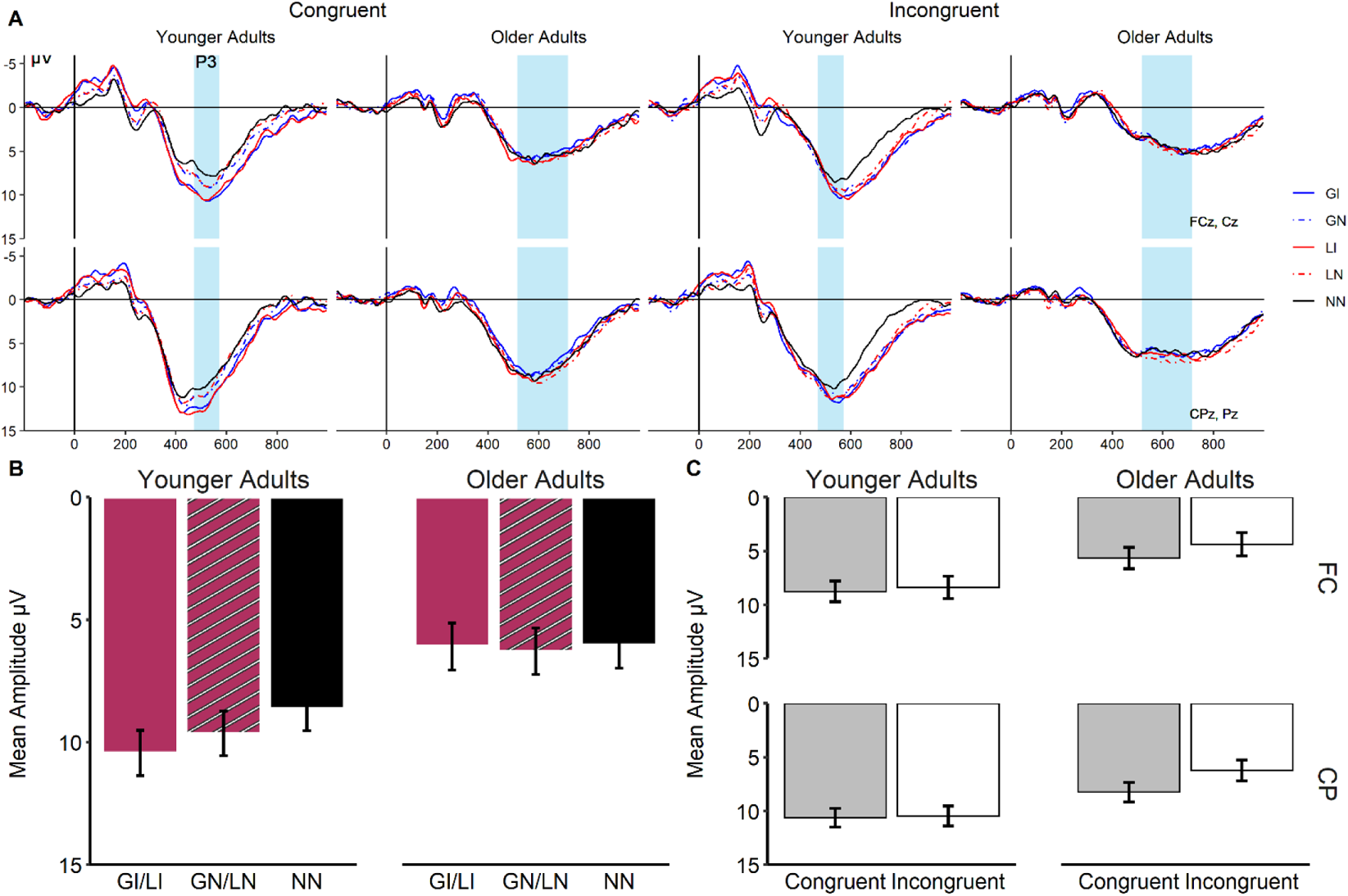
Target-Locked ERPs by Congruency, Age Group, Trial Type, and Region for P3 Amplitude *Note.* Shown in Panel A are ERP waveforms are averaged at frontocentral sites (top) and centroparietal sites (bottom). The blue shaded region corresponds to the time window for P3 amplitude. GI = gain-incentive trials; GN = gain-neutral trials; LI = loss-incentive trials; LN = loss-neutral trials; NN = neutral-neutral trials. Panel B represents the estimated marginal means from the age group by incentive interaction. Panel C represents the estimated marginal means from the age group by congruency by region interaction. FC (frontocentral), CP (centroparietal).

Second, a multilevel logistic regression model (random intercept only) was estimated with trials nested within subjects, regressing accuracy (i.e., incorrect = 0, correct = 1) onto age group (younger, older), congruency (congruent, incongruent), ANT cue (no, double), and trial type collapsed across valence (i.e., incentive trials [GI, LI], mixed-block neutral trials [GN, LN], and neutral trials [NN]). For both models we specified up to 2-way interactions of factors because comparisons with models that included higher-order interactions indicated that the additional parameters did not yield a better fit for the RT model, *Χ^2^* (9) = 5.37, *p* = .80, (AIC_Full_ = 272116, BIC_Full_ = 272324; AIC_Reduced_ = 272103, BIC_Reduced_ = 272239), or for the accuracy model, *Χ^2^* (9) = 5.74, *p* = .77, (AIC_Full_ = 9450.3, BIC_Full_ = 9652.5; AIC_Reduced_ = 9438.0, BIC_Reduced_ = 9567.4). Follow-up pairwise comparisons of interest for both behavioral and neurophysiological data were the transient (i.e., incentive vs. mixed-block neutral trials) and sustained (i.e., mixed-block neutral vs. neutral trials) effects of incentives.

### ERP Data Analysis

ERP analyses were performed in relation to the incentive cue, ANT cue, and target-evoked activity. For the incentive cue, mean amplitudes were averaged at frontocentral (FCz and Cz) and centroparietal (CPz and Pz) sites for the P2 and early negative component, and at frontocentral (FCz and Cz) sites for the CNV. Incentive cue-P2 mean amplitude was analyzed during the time window of 200-300ms. For the incentive cue early negative component analysis, we first extracted peak latencies based on the morphology of the waveform, using the time window of 300-500ms for younger adults and 350-600ms for older adults. Mean amplitudes were defined using a 100ms window centered on group averaged peak latencies. Mean amplitude of the early negative component was analyzed using a mixed-factorial ANOVA with age group (younger, older) as the between-subjects variable and region (frontocentral, centroparietal) and trial type (incentive, mixed-block neutral, neutral) as the within-subjects factors. CNV mean amplitude was examined 700-1000ms following the incentive cue. The mixed-factorial ANOVA for CNV activity was conducted with age group (younger, older) as the between-subjects variable and trial type (incentive, mixed-block neutral, neutral) as the single within-subjects factor.

Posterior N1 was examined 150-250ms following either (1) the alerting cue or (2) the continuation of the fixation that was on-screen at the time of the alerting cue in the no cue condition. Posterior N1 amplitude was averaged at both parietal (P3, Pz, and P4) and occipital (O1, Oz, and O2) sites. The mixed-factorial ANOVA for ANT cue-locked N1 mean amplitude was conducted with age group (younger, older) as the between-subjects variable and region (parietal, occipital), ANT cue type (no cue, double cue), and trial type (incentive, mixed-block neutral, neutral) as the within-subjects factors. In addition, posterior N1 was examined 150-250ms after the presentation of the target. As in the cue-locked analysis, average posterior N1 amplitude was analyzed at parietal (P3, Pz, and P4) and occipital (O1, Oz, and O2) electrodes. The target-locked posterior N1 analysis was conducted with age group (younger, older) as the between-subjects variable and region (parietal, occipital), congruency (congruent, incongruent), and trial type (incentive, mixed-block neutral, neutral) as the within-subjects factors.

For the analysis of target-locked P3 amplitude, we examined both frontocentral (FCz, Cz) sites and centroparietal (CPz, Pz). We first extracted peak latencies, using the time window of 300-800ms for both age groups following the presentation of the target. As Williams et al. (2016) demonstrated that the P3 component exhibits a wider distribution in older adults, mean amplitudes were calculated over a 200ms window for older adults, and a 100ms window for younger adults, centred at group averaged peak latencies. The mixed-factorial ANOVA for target-locked P3 mean amplitude was conducted with age group (younger, older) as the between-subjects variable and region (frontocentral, centroparietal), congruency (congruent, incongruent), and trial type (incentive, mixed-block neutral, neutral) as the within-subjects factors. In situations where Mauchly’s assumption of sphericity was violated, the Huynh-Feldt correction was used to adjust degrees of freedom. Benjamini-Hochberg adjusted *p*-values (Benjamini & Hochberg, 1995) were used to account for multiple comparisons.

## Results

### Behavioral Results

#### Correct RT

Means and standard deviations for behavioral measures are presented in Table 2 and the results from the analysis-of-variance table calculated on the multilevel linear regression model for RT as well as effect sizes are presented in Table 3. The model revealed participants were faster to respond on congruent vs. incongruent trials, and on double cue vs. no cue trials. There was also a main effect of trial type. Comparisons of interest revealed both transient, *p* < .001, *d* = .15, and sustained, *p* < .001, *d* = .16, effects of incentives, with faster responses on incentive vs. mixed-block neutral trials (i.e., transient) and faster responses on mixed-block neutral trials vs. neutral trials (i.e., sustained).

**Table 2.**
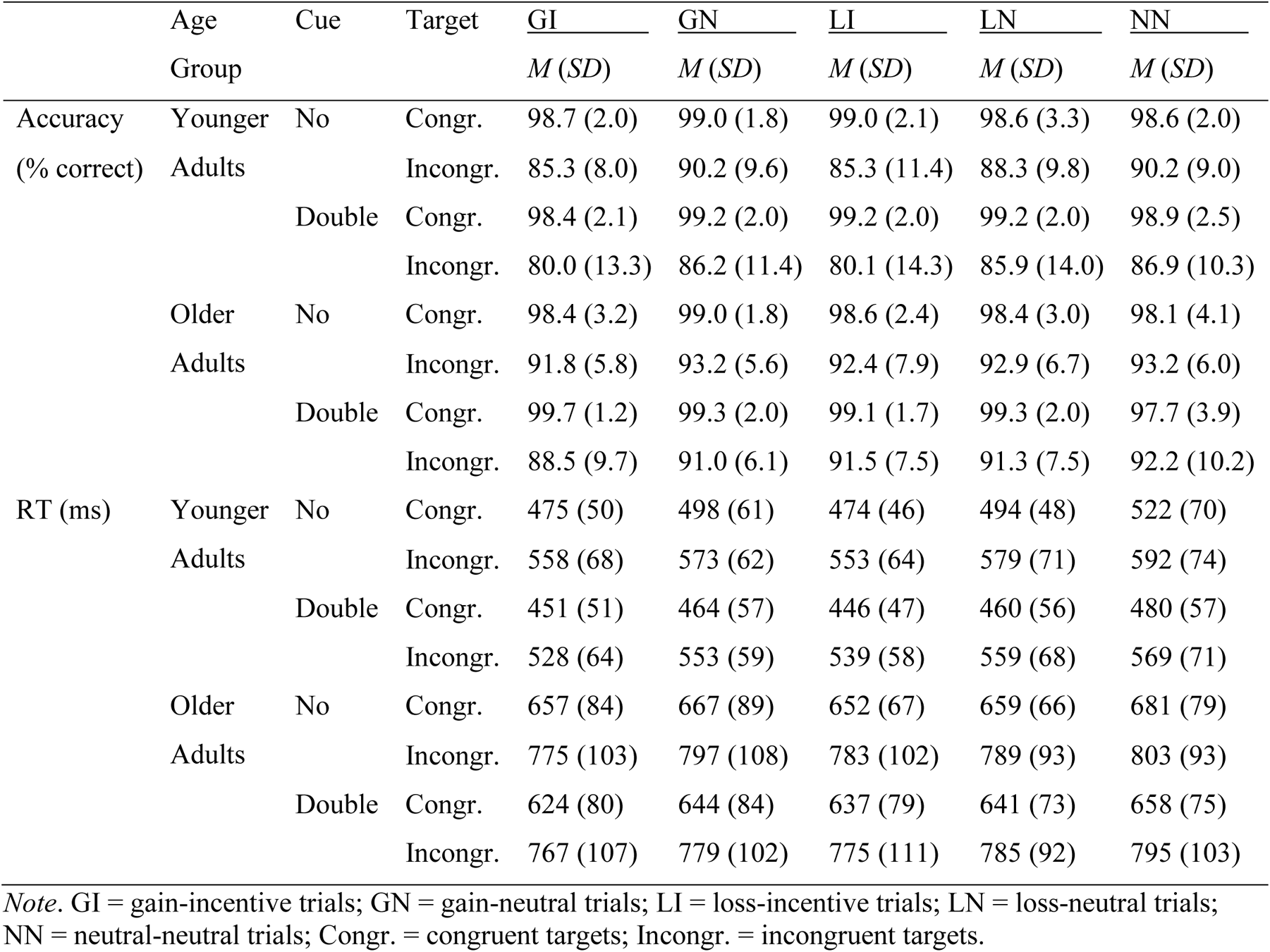
Means (M) and Standard Deviations (SD) for Behavioral Data

**Table 3.** Main Effects and Interactions for Correct RT

| Correct RT |  |  |  |  |
| --- | --- | --- | --- | --- |
| Analysis of Deviance Table (Type III Wald chi-square tests) |  |  |  |  |
| | $\chi^2$ | *Df* | *p* | $\eta_p^2$ |
| Intercept | 1257.42 | 1 | < .001 |  |
| Age Group | 69.25 | 1 | < .001 | .67 |
| Congruency | 815.02 | 1 | < .001 | .20 |
| Cue | 138.62 | 1 | < .001 | .01 |
| Trial Type | 134.11 | 2 | < .001 | .01 |
| Age Group x Congruency | 313.48 | 1 | < .001 | .01 |
| Age Group x Cue | 17.46 | 1 | < .001 | 7.77e-04 |
| Congruency x Cue | 19.94 | 1 | < .001 | 8.88e-04 |
| Age Group x Trial Type | 13.53 | 2 | .001 | 6.02e-04 |
| Congruency x Trial Type | 3.53 | 2 | .17 | 1.57e-04 |
| Cue x Trial Type | 1.00 | 2 | .61 | 4.45e-05 |

In addition, there was a main effect of age group, and age group interacted separately with trial type, congruency, and cue. Younger adults showed transient, *p* < .001, *d* = .19, and sustained speed-up effects, *p* < .001, *d* = .18, and older adults had transient, *p* < .001, *d* = .11, and sustained, *p* < .001, *d* = .14, speed-up effects, with both age groups exhibiting faster RT for incentive vs. mixed-block neutral trials and faster RT for mixed-block neutral trials vs. neutral trials. When further compared, the transient effect was larger in younger adults vs. older adults, *p* = .005, but there was no age difference for the sustained effect, *p* = .34. Both younger adults, *p* < .001, *d* = .83, and older adults, *p* < .001, *d* = 1.30, were faster on congruent vs. incongruent trials, with the difference being larger for older adults vs. younger adults, *p* < .001. Furthermore, both younger adults, *p* < .001, *d* = .27, and older adults, *p* < .001, *d* = .16, were faster on double cue vs. no cue trials, but this alerting effect was larger for younger adults vs. older adults, *p* < .001.

Finally, there was a significant interaction of congruency by cue. The presence of the double cue vs. no cue resulted in faster responses for both congruent, *p* < .001, *d* = .27, and incongruent trials, *p* < .001, *d* = .15, with a smaller alerting effect for incongruent vs. congruent trials, *p* < .001.

#### Accuracy

The results from the analysis-of-variance table calculated on the multilevel logistic regression model for accuracy are presented in Table 4. Results of the model indicated a significant main effect of congruency, which interacted separately with age group, cue, and trial type. There was no age group difference in accuracy on congruent trials, *p* = .61, *OR* = 0.88, but older adults were more likely to be accurate than younger adults on incongruent trials, *p* = .003, *OR* = 1.79. For congruent trials, there was no significant difference in accuracy with respect to the no cue vs. double cue condition, *p* = .08, *OR* = .74, whereas for incongruent trials, participants were more likely to be accurate in the no cue vs. double cue condition, *p* < .001, *OR* = 1.35. Accuracy was higher for all three trial types for congruent vs. incongruent trials, *p*s < .001, however this congruency effect was larger for mixed-block neutral trials compared to neutral trials (i.e., sustained effect), *p* = .01, *OR* = 1.83, with no congruency difference between mixed-block neutral trials and incentive trials (i.e., transient effect), *p* = .40, *OR* = .84.

**Table 4.**
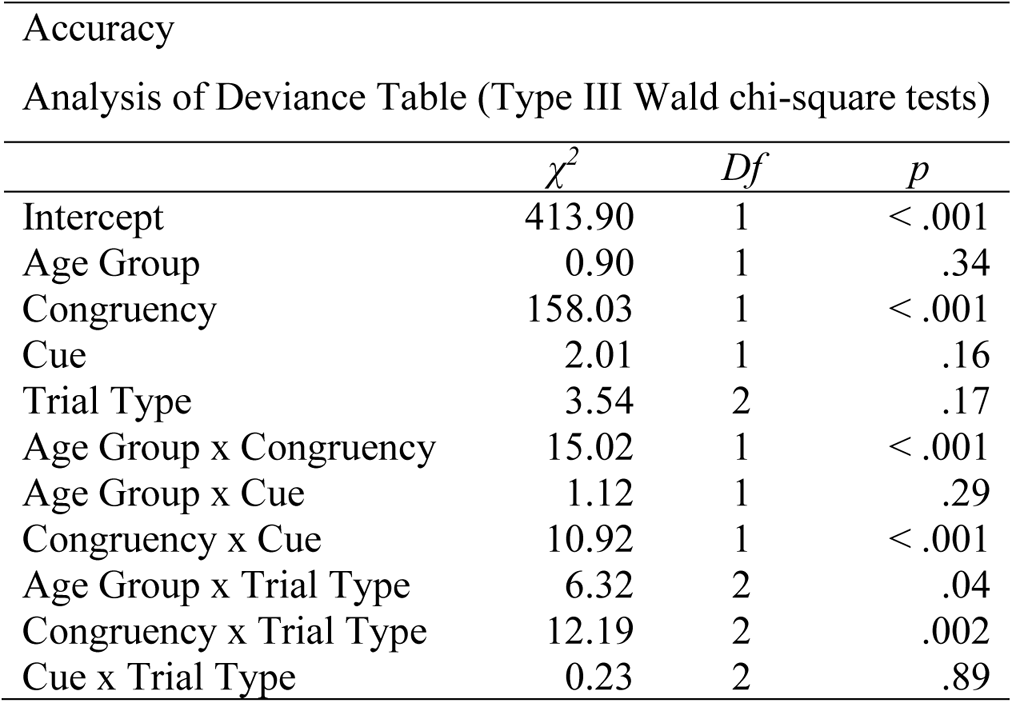
Main Effects and Interactions for Accuracy

Finally, the age group by trial type interaction revealed that older adults were more likely to be accurate than younger adults on incentive trials, *p* < .05, *OR* = 1.54, with no age group difference shown for mixed-block neutral trials, *p* = .46, *OR* = 1.17, or neutral trials, *p* = .71, *OR* = 1.09. Trial type comparisons revealed a significant transient effect for younger adults, *p* = .02, *OR* = 1.38, but not older adults, *p* = .65, *OR* = 1.06, with higher accuracy for mixed-block neutral vs. incentive trials. Sustained effects were nonsignificant for both younger adults, *p* = .28, *OR* = 1.20, and for older adults, *p* = .24, *OR* = 1.30.

#### Summary of Behavioral Results

The presence of incentives was associated with significant transient and sustained speed-up effects for both age groups, but the transient effect was larger for younger adults. Younger adults also exhibited a larger alerting effect for RT compared to older adults. Older adults slowed down more than younger adults on incongruent vs. congruent trials. Faster responding by younger adults was associated with reduced accuracy compared to older adults, for both incongruent trials and incentive trials (see Table 5 for a summary of incentive effects).

**Table 5.** Summary of Incentive Effects by Age Group

|  |  | <u>Younger Adults</u> |  | <u>Older Adults</u> |  |
| --- | --- | --- | --- | --- | --- |
|  |  | Transient | Sustained | Transient | Sustained |
| **Behavioral Measures** |  |  |  |  |  |
|  | RT | x<sup>a</sup> | x | x | x |
|  | Accuracy | x<sup>b</sup> | - | - | - |
| **ERP Components** |  |  |  |  |  |
| Incentive Cue |  |  |  |  |  |
|  | P2 | x | x | x | x |
|  | Early Negative | x | - | - | x |
|  | CNV | x | - | - | x |
| ANT Cue |  |  |  |  |  |
|  | N1 | x | x | x | x |
| Target |  |  |  |  |  |
|  | N1 | x | x | - | - |
|  | P3 | x | x | - | - |

x = Significant effect.
<sup>a</sup> Effect is larger in younger adults.<sup>b</sup> Accuracy cost on incentive trials.

### ERP Results

#### Incentive Cue-Locked Results

##### P2 Amplitude

Average ERP waveforms time-locked to the incentive cue are depicted in Figure 2 for each age group and region. For P2 mean amplitude, there were main effects of trial type, *F*(1.69, 81.28) = 14.31, *p* < .001, η_p_^2^ = .23, and region, *F*(1, 48) = 13.56, *p* < .001, η_p_^2^ = .22. Region interacted with age group, *F*(1, 48) = 16.22, *p* < .001, η_p_^2^ = .25, and trial type, *F*(1.84, 88.24) = 8.14, *p* < .001, η_p_^2^ = .15. Lastly, there was a three-way interaction between trial type, region, and age group, *F*(1.84, 88.24) = 3.34, *p* = .04, η_p_^2^ = .07. As shown in the yellow-shaded regions of Figure 2, younger adults showed transient and sustained effects of incentives at both frontocentral, *p*_BH_s < .001, and centroparietal regions, *p*_BHtransient_ < .01; *p*_BHsustained_ < .001, with larger P2 amplitude shown for incentive vs. mixed-block neutral trials, and for neutral vs. mixed-block neutral trials. Older adults showed the same P2 amplitude pattern for transient and sustained effects at frontocentral sites, *p*_BHtransient_ < .01; *p*_BHsustained_ = .01, but only a sustained effect was significant at centroparietal sites, *p*_BH_ < .05, while the transient effect failed to reach significance, *p*_BH_ = .09. There were no other significant effects or interactions, *F*s ≤ 1.22.

##### Early Negative Component Amplitude

For this component, there was a main effect of region, *F*(1, 48) = 42.94, *p* < .001, η_p_^2^ = .47, with more negative amplitude at frontocentral (*M* = −1.28, *SE* = 0.35) vs. centroparietal sites (*M* = −0.33, *SE* = 0.33). There was also a main effect of trial type, *F*(1.62, 77.73) = 6.83, *p* = .004, η_p_^2^ = .13, which interacted with age group, *F*(1.62, 77.73) = 4.83, *p* = .016, η_p_^2^ = .09. As shown in the blue-shaded regions of Figure 2, younger adults showed a significant transient effect of incentives, *p*_BH_ < .001, with reduced negative amplitude for incentive vs. mixed-block neutral trials, whereas older adults showed a significant sustained effect of incentives, *p*_BH_ < .05, with more negative amplitude for mixed-block neutral trials vs. neutral trials. The sustained effect of incentives was nonsignificant for younger adults, *p*_BH_ = .36, and the transient effect of incentives was nonsignificant for older adults, *p*_BH_ =.18. There were no other significant main effects or interactions, *F*s ≤ 2.58.

##### CNV Amplitude

There was a main effect of age group, *F*(1, 48) = 4.68, *p* = .035, η_p_^2^ = .09, trial type, *F*(2, 96) = 3.25, *p* = .043, η_p_^2^ = .06, and a significant age group by trial type interaction, *F*(2, 96) = 3.86, *p* = .024, η_p_^2^ = .07. As shown in the pink-shaded regions of Figure 2, younger adults displayed a significant transient effect of incentives, *p*_BH_ < .05, but no sustained effect, *p*_BH_ = .58, with more negative amplitude for incentive vs. mixed-block neutral trials. In contrast, older adults displayed a sustained effect of incentives, *p* = .03; *p*_BH_ = .06, but not a transient effect, *p*_BH_ =.51, with more negative amplitude for mixed-block neutral trials vs. neutral trials.

#### ANT Cue-Locked Results

##### ANT Cue-N1 Amplitude

Difference waves depicting the double cue minus the no cue condition (i.e., alerting effect) are presented in Figure 3. There was a main effect of age group, *F*(1, 48) = 17.02, *p* < .001, η_p_^2^ = .26, trial type, *F*(2, 96) = 13.97, *p* < .001, η_p_^2^ = .23, and region, *F*(1, 48) = 19.55, *p* < .001, η_p_^2^ = .29. Age group interacted with both cue type, *F*(1, 48) = 16.99, *p* < .001, η_p_^2^ = .26, and trial type, *F*(2, 96) = 12.92, *p* < .001, η_p_^2^ = .21. Younger adults (*M* = −1.67, *SE* = 0.36) had more negative amplitude than older adults (*M* = 0.57, *SE* = 0.37) for double cue trials, *p*_BH_ < .001, however there was no age difference in N1 amplitude for no cue trials, *p*_BH_ = .84 (*M_younger_* = −0.23, *SE_younger_* = 0.12; *M_older_* = −0.27, *SE_older_* = 0.12). Both younger and older adults showed transient, *p*_BHyounger_ < .001; *p*_BHolder_ < .05, and sustained, *p*_BH_s < .05, effects of incentives, irrespective of ANT cue type. Younger adults had more negative amplitude for incentive (*M* = −1.52, *SE* = 0.20) vs. mixed-block trials (*M* = −0.85, *SE* = 0.20), and for mixed-block trials vs. neutral trials (*M* = −0.47, *SE* = 0.21). Older adults showed reduced amplitude for incentive trials (*M* = 0.06, *SE* = 0.21) vs. mixed-block trials (*M* = 0.37, *SE* = 0.21), and for neutral trials (*M* = 0.02, *SE* = 0.22) vs. mixed-block trials.

There was also a region by trial type interaction, *F*(2, 96) = 6.29, *p* = .003, η_p_^2^ = .12. Transient effects of incentives were present at both parietal (*M*_Incentive_ = −0.58, *SE*_Incentive_ = 0.16_;_ *M*_MixedNeutral_ = 0.001, *SE*_MixedNeutral_ = 0.16), *p*_BH_ < .001, and occipital sites (*M*_Incentive_ = −0.88, *SE*_Incentive_ = 0.15_;_ *M*_MixedNeutral_ = −0.48, *SE*_MixedNeutral_ = 0.15), *p*_BH_ = .001, whereas there were no sustained effects at parietal (*M*_MixedNeutral_ = 0.001, *SE*_MixedNeutral_ = 0.16; *M*_Neutral_ = 0.07, *SE*_Neutral_ = 0.17) or occipital sites (*M*_MixedNeutral_ = −0.48, *SE*_MixedNeutral_ = 0.15; *M*_Neutral_ = −0.51, *SE*_Neutral_ = 0.16), *p*_BH_s = .79.

Finally, there was also a region by cue type interaction, *F*(1, 48) = 36.68, *p* < .001, η_p_^2^ = .43. In general, double cue trials (*M* = −1.07, *SE* = 0.27) had more negative amplitude than no cue trials (*M* = −0.18, *SE* = 0.08) at occipital sites, *p*_BH_ < .01, whereas there was no significant difference in amplitude between double cue (*M* = −0.02, *SE* = 0.28) and no cue (*M* = −0.32, *SE* = 0.10) trials at parietal sites, *p*_BH_ = .33. There were no other significant main effects or interactions, *F*s ≤ 1.23.

#### Target-Locked Results

##### Target-N1 Amplitude

For posterior N1 amplitude at the level of the target, there was a main effect of age, *F*(1, 48) = 5.85, *p* = .019, η_p_^2^ = .11, and a main effect of trial type, *F*(1.67, 80.29) = 16.45, *p* < .001, η_p_^2^ = .26. There was also a significant trial type by age group interaction, *F*(1.67, 80.29) = 9.07, *p* < .001, η_p_^2^ = .16, as well as a significant trial type by age group by region interaction, *F*(1.87, 89.74) = 3.58, *p* = .035, η_p_^2^ = .07. As shown in the blue-shaded region of Figure 4, for younger adults, there were both transient, *p*_BH_s < .001, and sustained, *p*_BH_s < .05, effects of incentives at parietal and occipital sites, whereas there were no significant incentive effects for older adults at either site, *p*_BH_s ≥ .22. Younger adults had larger N1 amplitude for incentive vs. mixed-block neutral trials and for mixed-block neutral vs. neutral trials. There were no other significant main effects or interactions, *F*s ≤ 2.97.

##### Target-P3 Amplitude

Younger adults had larger target-locked P3 amplitude compared to older adults, *F*(1, 48) = 7.35, *p* = .009, η_p_^2^ = .13. Larger P3 amplitude was also present for congruent vs. incongruent trials, *F*(1, 48) = 5.71, *p* = .021, η_p_^2^ = .11, and for centroparietal sites vs. frontocentral sites, *F*(1, 48) = 33.58, *p* < .001, η_p_^2^ = .41. There was also a main effect of trial type, *F*(1.75, 84.11) = 4.69, *p* = .015, η_p_^2^ = .09, and trial type interacted with age group, *F*(1.75, 84.11) = 3.96, *p* = .027, η_p_^2^ = .08. As shown in the blue-shaded regions of Figure 5A and in the bar graph presented in Figure 5B, younger adults showed transient, *p* < .05; *p*_BH_ = .086, and sustained effects, *p*_BH_ < .05, of incentives, whereas there were no incentive effects for older adults, *p*_BH_s = .60. Younger adult P3 amplitude was larger for incentive vs. mixed-block neutral trials, and for mixed-block neutral vs. neutral trials.

There was also a significant three-way interaction of age group, congruency, and region, *F*(1, 48) = 8.75, *p* = .005, η_p_^2^ = .15. As shown in Figure 5C, younger adults showed no effect of congruency at either frontocentral, *p*_BH_ = .71, or centroparietal sites, *p*_BH_ = .77. Older adults demonstrated an effect of congruency, which was larger at centroparietal, *p*_BH_ < .01, vs. frontocentral sites, *p* < .05; *p*_BH_ = .07. Finally, there was a significant interaction of trial type by congruency by region, *F*(1.81, 87.03) = 3.42, *p* = .042, η_p_^2^ = .07. There was a significant congruency effect for incentive trials at frontocentral and centroparietal sites, *p*_BH_s < .05, mixed-block neutral trials did not show any congruency effects, *p*_BH_s ≥ .26, whereas neutral trials only showed a significant congruency effect at centroparietal sites, *p*_BH_ < .05, but not frontocentral sites, *p*_BH_ = .31. There were no other significant main effect or interactions, *F*s ≤ 3.02.

#### Summary of ERP Results

A summary of the incentive effects for each age group, for behavioral and ERP measures, is presented in Table 5. For incentive cue-P2 amplitude, younger and older adults displayed a similar pattern of transient and sustained effects of incentives. Age differences in incentive processing emerged at the time of the early negative component and the subsequent CNV. Here, only transient effects were found for younger adults, whereas only sustained effects were present for older adults. For ANT cue-N1 amplitude, younger adults exhibited a stronger alerting effect than older adults. Both age groups showed transient and sustained effects of incentives, which were not influenced by the presence of the ANT cue. In term of target-locked N1 amplitude, only younger adults showed transient and sustained effects of incentives at this stage of processing.

Similar effects were present for target-locked P3 amplitude, with transient and sustained effects present for younger adults, but not older adults. Finally, only older adults showed a significant congruency effect, with smaller P3 amplitude for incongruent vs. congruent trials. This effect was larger at centroparietal vs. frontocentral sites.

## Discussion

The current study sheds new light on the time course of incentive effects on attention in younger and older adults. By examining early and late occurring ERP components within the course of the trial, as well as the transient vs. sustained nature of these effects, we characterized age differences in the temporal dynamics of incentive processing across multiple timescales. Incentives were associated with both transient and sustained speed-up effects on RTs, with larger transient effects shown in younger adults. Alerting cues had a larger impact on younger adults’ response times, whereas older adults showed larger congruency effects. Younger adults were less accurate than older adults on incentive trials and incongruent trials. ERP analyses at the time of the incentive cue showed the same pattern of transient and sustained effects of incentives for P2 amplitude across age groups. Following the P2, younger adults showed transient effects whereas older adults showed sustained effects of incentives for the early negative component and CNV. At the time of the alerting cue, N1 amplitude was larger for younger adults, and showed transient and sustained incentive-related increases in younger adults, but decreases in older adults. At the time of the target, younger adults had larger N1 and P3 amplitudes and continued to show transient and sustained incentive effects, whereas older adults did not show any effects of incentives. We consider each of these results in turn with respect to alterations in dopaminergic neuromodulation in aging.

### Incentive- and alerting-induced speed-up effects reflect age differences in preparatory activity and response strategies

Consistent with Williams et al. (2018), incentives were associated with both transient and sustained speed-up effects. As the transient RT effect was larger for younger adults, it appears that incentives were more effective at amplifying trial-to-trial level preparatory activity and updating processes for younger vs. older adults. Use of MLMs in our behavioral analyses allowed for increased statistical power to assess trial-level effects as opposed to coarser averaging across trials using mixed-factorial ANOVAs (Quené & van den Bergh, 2004).

Inclusion of trial-level incentive cues as opposed to block-level incentive information was also more conducive to transient effects (Kostandyan et al., 2019). Enhanced preparatory activity in younger adults was also present via a larger alerting effect on RT in relation to older adults, and corresponds to typical age differences seen in prior literature (e.g., Gamboz et al., 2010; Jennings et al., 2007). Considering that incentive and alerting effects did not interact, this suggests that the different cue types may be eliciting additive rather than interactive effects on preparatory processes.

As older adults were slower and had higher accuracy on incongruent and incentive trials compared with younger adults, these findings replicate previous reports of age differences in speed-accuracy trade-off settings (e.g., Williams et al., 2018). Incentive cues and alerting cues did not reduce flanker interference in either age group. It is possible that incentive-driven upregulation of cognitive control is limited to situations in which cues predict target identity or location (e.g., Chiew & Braver, 2016).

### Age differences in transient vs. sustained incentive effects emerge at later stages of preparatory control

Similar to Schmitt et al. (2015), at the time of the incentive cue both age groups showed transient increases in P2 amplitude for incentive vs. neutral trials. The transient effect for older adults was more pronounced at frontocentral relative to centroparietal sites whereas younger adults had transient effects at both sites, which may reflect greater reliance on frontal attentional resources by older adults. Sustained effects were present for both age groups at both sites. As amplitude was lower for neutral trials embedded in incentive blocks vs. pure blocks, this suggests that in the mixed-block context greater allocation of attention is directed towards incentive cues vs. neutral cues. Importantly, these results indicate that the early, automatic attentional processes associated with incentive cues are preserved with age.

Following the P2, we focused on an early negative component observed during a similar time window as the incentive cue-locked P3 component in Williams et al. (2018). It was at this stage of processing that age differences emerged in the form of a dissociation, whereby younger adults showed transient incentive effects and older adults displayed sustained effects. Due to the negative morphonology of this component, we suspect this may be an N450 effect. Analogous to the P3, the N450 is involved in cognitive as well as emotional conflict monitoring and detection (Ma, Liu, Zhong, Wang, & Chen, 2014; Zhu et al., 2018). As the transient effect in younger adults was represented by attenuated negative amplitude for incentives in relation to mixed-block neutral trials, this may represent more efficient processing of incentives by younger adults at this stage in the trial.

For older adults, the presence of a sustained but not transient effect suggests that older adults processed incentives and neutral trials in a similar manner within the mixed-block context. The presence of an N450 component rather than a P3 component during this time window may be due to shorter incentive cue presentation (200ms vs. 400ms) than Williams et al. (2018) or may be due to participants experiencing a different anticipatory context with the addition of the alerting cue. Consequently, while Williams et al. (2018) found sustained effects at the time of the incentive cue-P3 for both age groups, there is a lack of a sustained effect for younger adults in the current study at the time of this early negative component. However, we note that around 600ms for younger adults in the current study there is, visually, a graded positive effect that is similar to what was shown for the P3 component for younger adults in the Williams et al. (2018) study. Therefore, there may be a shift in the time course due to the addition of the alerting cue.

The age-related dissociation continued late into incentive cue processing during the time window of the CNV component. These results also support greater anticipatory activity by younger adults with larger CNV amplitude for incentive trials, whereas older adults continue to show similar preparatory activity for incentive and neutral trials within the mixed-block context. These results differ from Williams et al. (2018), in which both age groups displayed transient effects of incentives for CNV amplitude, but aligns with Schmitt et al. (2015) whereby younger adults displayed larger CNV amplitude for loss incentive cues in a design that incorporated a secondary context cue following the incentive cue. Thus, in a dual-cue context older adults may resort to response strategies that are more likely associated with sustained preparatory activity to deal with increased processing demands from the combination of incentive and alerting cues.

Following the incentive cue, younger adults had a more prominent alerting effect via N1 amplitude than older adults. Williams et al. (2016) found no age differences for ANT cue-locked N1 when incentive information was presented at the block-level, and therefore the current study’s trial-level anticipatory incentive cue context may indirectly exacerbate age differences in the alerting network. Although both age groups showed transient and sustained effects of incentives, younger adults showed a more predictable, graded negative N1 amplitude pattern that scaled with incentives, whereas older adults had a more attenuated effect. Consequently, at the time point of the alerting cue incentive effects became more fleeting for older adults compared to younger adults.

### Transient and sustained effects of incentives are present during target processing for younger but not older adults and do not enhance cognitive control

In support of younger adults showing an enhanced state of preparatory attention throughout the trial, younger adults also had larger N1 amplitude than older adults at the time of the target and showed transient and sustained effects of incentives, whereas at this time point incentive effects were absent for older adults. Replicating Williams et al. (2018), younger adults had larger target-P3 amplitude than older adults. Transient and sustained incentive effects for P3 amplitude were again present only for younger adults. Hence, incentives continued to influence early attentional processes (N1) at the time of the target and later occurring conflict detection processes (P3) for younger adults only. Similar to Williams et al. (2016; 2018), older adults showed a congruency effect for P3 amplitude, whereas younger adults exhibited similar amplitudes and processing resources for both trial types. Older adults showed declines in P3 amplitude for incongruent trials, but compensated via increased response times. While incentive effects were present for younger adults during target processing, incentives did not improve flanker interference and were instead associated with increased errors. Therefore, the heightened readiness in younger adults associated with incentives and alerting may be hard to counteract when conflict resolution is required at the time of the target.

### Implications for age differences in dopamine transmission

The results of the current study provide greater insight into age differences in the temporal dynamics of incentive processing and may reflect age-related alterations across varying timescales of dopamine transmission. Early on at the incentive cue both age group show transient and sustained effects of incentives for the P2 component, which may implicate a dual role of rapid and continual dopamine transmission in preserving automatic attention with age. A dissociation emerged at mid-to-late stages of incentive cue processing via N450 and CNV amplitude. As younger adults solely showed transient effects, this suggests that more rapid dopamine transmission may be associated with enhanced trial-to-trial preparatory activity. The shift in older adults towards sustained, block-level processing may be a compensatory mechanism for age-related declines in transient dopamine transmission. A second modification took place at the time of the alerting cue. Younger adults continued to show rapid, transient effects of incentives for N1 amplitude, but also showed sustained effects that may have built up over time within the trial from gradual dopamine levels. Although transient and sustained effects were present for older adults at the time of the alerting cue, they were largely diminished due to a reduced N1 response. Furthermore, at the target younger adults maintained transient and sustained effects for N1 and P3 amplitude, whereas incentive effects were absent in older adults and may reflect overall declines in dopamine transmission with age during target processing.

## Conclusion

Early automatic attentional processes associated with incentives were found to be preserved in older age. Age differences in the temporal dynamics of incentive processing emerged at later stages of the incentive cue, revealing a shift towards transient effects in younger adults vs. sustained effects in older adults and may correspond to age-related fluctuations in rapid vs. gradual dopamine release. Our results support enhanced preparatory processing with incentives in younger adults by means of transient effects at the incentive cue, which are followed by a ramping up of sustained effects to accompany transient effects at the time of alerting cue. Incentive effects in older adults were more fleeting and began to diminish at the alerting cue and were absent by the time of the target. Transient and sustained effects of incentives were maintained during target processing for younger adults. Future research should investigate whether the same temporal dynamics persist when predictive task cues are included (e.g., Chiew & Braver, 2016), and if such a context is more conducive to cognitive control benefits across age groups. Importantly, the current study provides novel evidence of when in the time course of incentive processing age difference arise and also the precise nature of these changes.

## CRediT Authorship Contribution Statement

**Margot D. Sullivan:** Formal analysis, Visualization, Writing - original draft, Writing - review & editing. **Farrah Kudus:** Conceptualization, Project administration, Investigation, Resources, Methodology, Formal analysis, Writing - original draft. **Benjamin J. Dyson:** Conceptualization, Writing - review & editing. **Julia Spaniol:** Supervision, Project administration, Conceptualization, Methodology, Writing - review & editing, Funding acquisition.

## Data Availability Statement

Due to institutional restrictions the raw data cannot be made publicly available. Data may be made available upon request to the corresponding author pending a formal data sharing agreement and ethics approval.

## Funding Information and Acknowledgments

This research was supported by the Canada Research Chairs Program (#950-232332 to J.S) and by the Natural Sciences and Engineering Research Council of Canada (RGPIN-2018-04455 to J.S.). We thank Carson Pun for his assistance with the study.

## Notes

1 Upon visual inspection, the ERP component elicited by the incentive cue in the current study, which corresponded to the time window of the incentive cue-P3 component in Williams et al. (2018), was negative rather than positive.

2 When valence was included, both age groups showed transient (gains only) and sustained (gains and losses) speed-ups of responses. However, the transient effect on RT for losses was only significant for younger adults, *p* < .001, and was marginal for older adults, *p* = .05.

